# Targeting ABCB4 using mRNA-LNP for the treatment of rare liver diseases

**DOI:** 10.1101/2023.04.11.535868

**Authors:** Mohammed Alsuraih, Brianna LaViolette, Guan-Yu Lin, Ramesh Kovi, Natalie Daurio, Congsheng Cheng, Youngwook Ahn, Zhihua Jiang, Roberto Ortiz, Shangzhong Li, Yuxing Cheng, Ye Wang, Xiaoyu Fan, Jessica Haskins, Xiuhua Sun, Abigail Hunter, Dinesh Hirenallur Shanthapa, Ying Wu, Matthew Holsti, Morag Stewart, Marija Tadin-Strapps, Shian-Huey Chiang

## Abstract

Mutations in the ABCB4 gene lead to a wide-spectrum of rare liver diseases including progressive familial intrahepatic cholestasis type 3 (PFIC3) and low-phospholipid associated cholelithiasis (LPAC) syndrome. PFIC3 patients develop symptoms during late infancy, including severe itching, jaundice, and failure to thrive. The condition may progress to liver failure during childhood or adulthood. This is a highly unmet medical condition where liver transplantation is the only option to correct this disease. Recently, exciting data suggested that restoration of the ABCB4 function via gene replacement could rescue liver phenotypes associated with ABCB4 dysfunction in a preclinical PFIC3 mouse model. Here, we used mRNA LNP platform to determine expression and durability of ABCB4 in the liver of wildtype mice. In addition, we generated Abcb4^-/-^ mice to study the efficacy of systemic delivery of ABCB4 mRNA LNP. We observed a robust and durable expression of hABCB4 up to 72 hours post systemic dosing in the liver of wild-type mice. Systemic administration of hABCB4 mRNA achieved a remarkable restoration of phosphatidylcholine levels in bile, a significant decrease in liver stiffness as measured by shear wave elastography, and amelioration of liver histopathology including fibrosis and ductular reaction. We conclude that administration of hABCB4 mRNA LNPs was sufficient to ameliorate fibrosis markers in the PFIC3 mouse model. Our data suggests that gene replacement using mRNA LNP modality could provide an excellent opportunity for patients with biliary diseases.

## INTRODUCTION

The multidrug resistance 3 (Mdr3) protein, which is encoded by the ATP-binding cassette (ABC) subfamily B member 4 (ABCB4) gene, is a member of the superfamily of ABC transporters [1, 2]. ABCB4 is an energy-dependent floppase, translocating the biliary phosphatidylcholine (PC) from the inner to the outer leaflet of the canalicular hepatocytes to form mainly PC-cholesterol vesicles and some mixed bile salt micelles. This mechanism protects the hepatobiliary system from deleterious detergent and lithogenic properties of the bile. Defects in this protein have been studied and functionally characterized, with multiple types of mutations resulting in low levels of biliary PC and destabilized micelles, as well as toxic bile leading to damage to the biliary epithelial cells and hepatocytes [3, 4].

A wide-spectrum of rare liver diseases is caused by mutations in the ABCB4 gene including progressive familial intrahepatic cholestasis type 3 (PFIC3), low-phospholipid associated cholelithiasis (LPAC) syndrome, and other rare biliary diseases (e.g., primary biliary cirrhosis) [5-9]. Moreover, variants in this transporter could play a role in driving liver fibrosis in patients with chronic liver diseases [10]. PFIC3 patients develop symptoms during late infancy, including severe itching, jaundice, and failure to thrive, and can progress to liver failure during childhood or adulthood. This is a medical condition with high unmet need with liver transplantation being the only option to correct this disease.

After the success of Covid-19 vaccines, mRNA has emerged as a potential therapeutic for treatment of both rare and common diseases. One drawback to this technology is the short-lived expression of the delivered mRNA, and consequently, the protein. Efforts are ongoing to develop more durable mRNA expression technologies, including development of circular mRNA and self-amplifying mRNA. Lipid nanoparticles (LNPs) delivery to organs other than the liver creates an additional challenge to expanding application to diseases beyond the liver. Indeed, rare liver diseases defined by monogenic mutations, present an opportunity to de-risk and demonstrate the promise of mRNA LNP therapeutics. To this end, a recent study demonstrated that delivery of ABCB4, using AAV or mRNA LNP platforms, could rescue the liver phenotypes associated with ABCB4 dysfunctions in a pre-clinical PFIC3 and primary sclerosing cholangitis (PSC) mouse model [11-13]. In an unrelated study, restoration of HNF4a via administration of HNF4a mRNA using LNPs inhibited liver fibrosis in both genetic and chemically induced pre-clinical models of liver fibrosis [14].

Therefore, we sought to further characterize the expression and durability of hABCB4 mRNA LNP *in vivo* and determine the efficacy of hABCB4 mRNA LNP in ameliorating liver stiffness, restoring healthy PC levels in bile, and correcting other phenotypes in the Abcb4^-/-^ mouse model of PFIC3 & PSC. Systemic administration of hABCB4 mRNA LNP resulted in robust Abcb4 protein expression *in vitro* and *in vivo*, which lasted for 72 hours, and consequently, led to a full restoration of PC level in bile and amelioration of liver disease phenotypes associated with the Abcb4^-/-^ mice.

## METHODS

### Genome-wide association study

The summary statistics of the GWAS meta-analysis were downloaded from https://www.med.umich.edu/spelioteslab/. The regional association plots for ±500kb window of the gene ABCB4 were created by using LocusZoom (http://locuszoom.org/). We used the LocusZoom-implemented LD estimates from 1000 Genomes Project (March 2012, EUR). The SNP rs2019505 (MAF 0.20) was applied as the index SNP and colored in purple in the plots.

### Animal work

All animal-involved procedures were reviewed and approved by Pfizer’s Institutional Animal Care and Use Committee (IACUC) and conducted in an AAALAC (Assessment and Accreditation of Laboratory Animal Care) International accredited facility. Mice deficient for the *Abcb4* gene were created and maintained on the FVB/NJ (JAX, #001800) strain, utilizing the CRISPR/Cas9 technology [29]. Knockout of *Abcb4* was achieved by introducing frame-shifting small insertions or deletions (Indels) into exon 7 of the gene. Cas9 guides were designed for efficient editing while minimizing potential risk of off-target editing (www.benchling.com). Seed sequences of Cas9 sgRNAs used for editing were; 5’GCTGATGGCCATGATCACGA (#696) and 5’ GGCTGATGGCCATGATCACG (#697). To generate F0 mice, fertilized embryos were harvested from super-ovulated females, electroporated with one of the two sgRNAs complexed with Cas9 protein (IDT) and implanted into pseudo-pregnant recipients. PCR genotyping of F0 pups was performed with genomic DNA extracted from tail biopsy using two primers, 5’CATGTGCACAGTGTGGATGC and 5’TTGGATGGCTGTGCTTTCAAC, followed by Sanger sequencing. Raw sequencing data were analyzed with the TIDE software (https://tide.nki.nl/) in order to measure frequency of individual Indels in each animal.

### mRNA-LNP generation

Messenger RNA substituted with N1-methylpseudouridine was synthesized *in vitro* from a DNA plasmid template. The template encoded a codon optimized full-length human ABCB4 (NCBI Reference Sequence: NP_000434.1). The template also provided 5’ and 3’ untranslated sequences and a poly-A tail. The mature RNA product had an N7-methylated cap to improve translation efficiency [30]. The final product was diluted in water prior to formulation into lipid nanoparticles (LNPs). **LNP formulations** were prepared using microfluidic mixing. In brief, lipids were dissolved in ethanol at molar ratios of 46.3:42.7:9.4:1.6 (ALC-0315:cholesterol: DSPC:ALC0159) as previously described [31, 32]. The lipid mixture was combined with a 50 mM citrate buffer (pH 4.0) containing mRNA at a ratio of 3:1 (aqueous: ethanol) using Precision Ignite NanoAssemblr (Precision Nanosystems, Vancouver, BC). LNPs were further placed into a 10K MWCO (molecular weight cut off) dialysis cassette and dialyzed in 10 mM Tris (pH=7.5) buffer overnight at 4 °C. The resulting formulations were concentrated using Amicon Ultra Centrifugal Filter devices 30KDa (EMD Millipore, Billerica, MA), sterile filtered using 0.22 µm PES filters (Fisher Scientific, NY, USA)) and 300 mM sucrose was added. All LNPs were tested for particle size, RNA encapsulation, RNA integrity and endotoxin (< 1 EU/ml of endotoxin).

### Phosphatidylcholine (PC) assay LC-MS

Lipids were extracted from bile samples using a liquidliquid extraction. Briefly, 600uL of ice cold 2:1 chloroform/methanol was added to 2 µL of bile. 2 µL of Lipidomix Splash Mix (Avanti Polar Lipids 330707) was then added to each sample and vortexed. 150 µL water was added to each sample and vortexed for approximately 20 seconds. Samples were incubated at room temperature for 5 minutes then spun at 10,000g for 10 minutes at 4C. Samples were then placed on ice and the bottom layer was transferred to a new tube and dried under a stream of N2. Residue was resuspended in 10 µL chloroform followed by 150 µL of methanol and transferred to a low volume glass HPLC vial. LC-IMS-MS analysis was performed on a Thermo Vanquish LC system coupled to a Bruker timsTOF pro mass spectrometer equipped with a VIP-HESI source. LC separation was performed on a YMC C18 2.1 × 100mm column at 55C. Mobile phase A was 10mM ammonium formate in 60% Acetonitrile and 40% water with 0.1% formic acid and mobile phase B was 10mM ammonium formate in 90% isopropanol 10% water with 0.1% formic acid. Gradient separation begun at 40% B and held for 2 minutes at a flow rate of 0.4mL/min. Gradient then increased to 50% B at 2 minutes and increased to 54% B over 10 minutes. Then increased to 70% B and ramped to 99% B over the next 6 minutes, held at 99%B for 2 minutes and returned to 40% for 2 minutes. Peak picking, integration and lipid annotation was performed by Bruker Metaboscape software. Bile PC concentrations were determined relative to the Splash Mix internal standard by lipid class.

### Protein expression LC-MS

Mouse liver was lysed and homogenized using 5% SDS (Fisher, BP2436200) in RIPA buffer (Sigma, R0278) with protease inhibitors (Thermo, PI87786), and the protein concentration of the lysate was measured using a BCA Protein Assay Kit (Thermo, 23225). 300 ug of total protein was processed and tryptic peptides were generated using the S-trap sample processing technology (Protifi). 5 fmol of stable isotope labeled internal standard (SIL) was added to the eluant. The eluant was then dried and resuspended in 250 µL 25mM Ammonium Formate (Sigma) and 100 µL was injected for IP-LCMS analysis tracking signature peptides for human and mouse ABCB4. Protein concentrations were calculated using the proportion of SIL, normalized to total protein of the lysate. The linear range was from 1 to 18 ng ABCB4/mg of total protein.

### Droplet Digital PCR Gene Expression Assay

A ddPCR assay was used for ABCB4. The reactions contained 2X Supermix, 20X ABCB4_FAM primer and probe mix (final concentration 450 and 200 nM, respectively), 0.04 U AmpErase UNG (Thermo Fisher), and 6.25 µL sample, QC, or water, in a total reaction volume of 25 µL. Droplets were generated in the Automated Droplet Generator (Bio-Rad). The plate was heat sealed with foil and droplets were amplified in a conventional thermal cycler with ramp rate 2.5°C per second (50°C for 2 minutes, 95°C for 10 minutes, 40 cycles of 94°C for 30 seconds and 60°C for 1 minute, followed by 98°C for 10 minutes). The fluorescence was measured on the QX200 Droplet Reader and the absolute nucleic acid copy count was processed on QX Manager software. Conversion of copy number to copies per µg total RNA was calculated in Excel.

### Shear wave Elastography Imaging

We adopted a shear wave elastography (SWE) imaging of liver from previously reported paper for measuring liver stiffness in mouse (Ross T et al, 2020; Morin J et al 2021). Briefly, SWE was performed using an Aixplorer Ultimate® system (SuperSonic Imagine, Aix-en-Provence, France) with 6 to 20 MHz linear array (SuperLinear™ SLH20-6) transducer. Mice were fasted for minimum of 4 hours to minimize interference of gut contents and movement during image acquisition. After fasting, mice were anesthetized using isoflurane (1 – 5%) and hair removed using Nair hair removal cream around the right lateral flank region for SWE imaging. After proper preparation of imaging area, a mouse was moved to a warmed surface, and placed in left lateral recumbency. After identifying clear area of liver on B-mode image of SWE, shear elasticity map of the tissue was generated using high-frequency ultrasound. Then, in built system tool was used to compute elasticity from a region of interest on shear wave elasticity map. Total of 3 to 5 shear wave mapping were captured from a randomly selected sections of liver area from an animal at each time point. Average elasticity value from these multiple measurements was used to report liver elasticity of a particular animal.

### Histology

FFPE sections were deparaffinized before incubation in Picro Sirius Red solution (Direct Red 81 0.2 % in 1.3 % Picric Acid Solution, Sigma). Sections were rinsed in dH2O, dried, mounted with Permount (Fisher) and acquired on a Leica system. Red pixels were quantified as percent area of fibrosis.

### Immunohistochemistry

Immunohistochemistry staining was performed in paraffin-embedded tissue sections. Briefly, 5µm paraffin sections were deparaffinized and re-hydrated, and slides were incubated with rabbit anti-human ABCB4 monoclonal antibody (1:8000, Clone: EPR23697-35, Abcam) antibody at overnight at 4°C. Immunohistological reaction of Abcb4 was developed using the peroxidase substrate ImmPACT DAB (Vector laboratories), and nuclei were counterstained with hematoxylin.

### Hydroxyproline quantification

100 mg of liver tissue were hydrolyzed in 1 mL of 6 N HCl at 110°C over night. 10 µL of the hydrolyzed sample or standard were placed in 30 µL of citric acetate buffer (10 g citric acid (5% w/v), 2.4 ml Glacial Acetic Acid (1.2% v/v), 14.48 g sodium acetate (7.24% w/v), 6.8 g sodium hydroxide (3.4% w/v) with a total volume of 200 ml made up with sterile deionized water). 100 µL of Chloramine T solution (0.282 g Chloramine T, 2 ml isopropanol, 2 ml sterile water, 16 ml citrate acetate buffer) were mixed with the samples or standards and oxidized for 20 min at room temperature. Oxidized samples or standards received 100 µL of Ehrlich’s Reagent (2.5 g of p-dimethylaminobenzaldehyde, 9.3 ml isopropanol, and 3.9 ml 70%-perchloric acid) and incubated at 65°C for 20 mins. Optical density at 550 nm was recorded and compared to the standard curve for quantification.

### Statistical analysis

All data are reported as the mean (or fold change in mean) ± SD from a minimum of 10 independent animals. Prism software was used for the generation of all bar graphs and statistical analyses. Statistical analyses were performed with 1way ANOVA, including Tukey’s multiple comparisons test for all comparisons between different animal groups.

## RESULTS

### Generation and Characterization of hABCB4 mRNA LNP *in vitro*

We generated two ABCB4 constructs; a wild-type full length, and S346I mutant human ABCB4. The S346I mutation affects the ability of the protein to translocate PC across the membrane without altering its expression level [7]. We transfected these two vectors into HEK293T and HuH7 cell lines and confirmed expression of hABCB4 (data not shown). We then generated wild-type, S346I mutant, and luciferase control modRNA by *in vitro* transcription, and tested for cell surface expression. hABCB4 expression in HuH7 cells was significantly increased (40% after 18 and 24 hours) over luciferase control by fluorescence-activated cell sorting (FACS) analysis (Fig. 1A). After successfully confirming the sequence and the expression of these constructs, wild-type hABCB4 and luciferase control were formulated into lipid nanoparticles (LNPs). We tested the expression of hABCB4 mRNA LNP in HuH7 cell line and observed an increase in hABCB4 protein level (Fig. 1B). In addition to cell surface expression, we tested the functionality of the hABCB4 mRNA LNP by measuring the level of PC in conditioned media. Huh7 cells were transfected with hABCB4 mRNA LNP and supernatants were then harvested at 24 hr post transfection for PC analysis using a commercially available kit. PC levels in the conditioned media were significantly increased in a dose-dependent manner upon hABCB4 mRNA LNP transfection relative to control cells (Fig. 1C).

**Figure 1.**
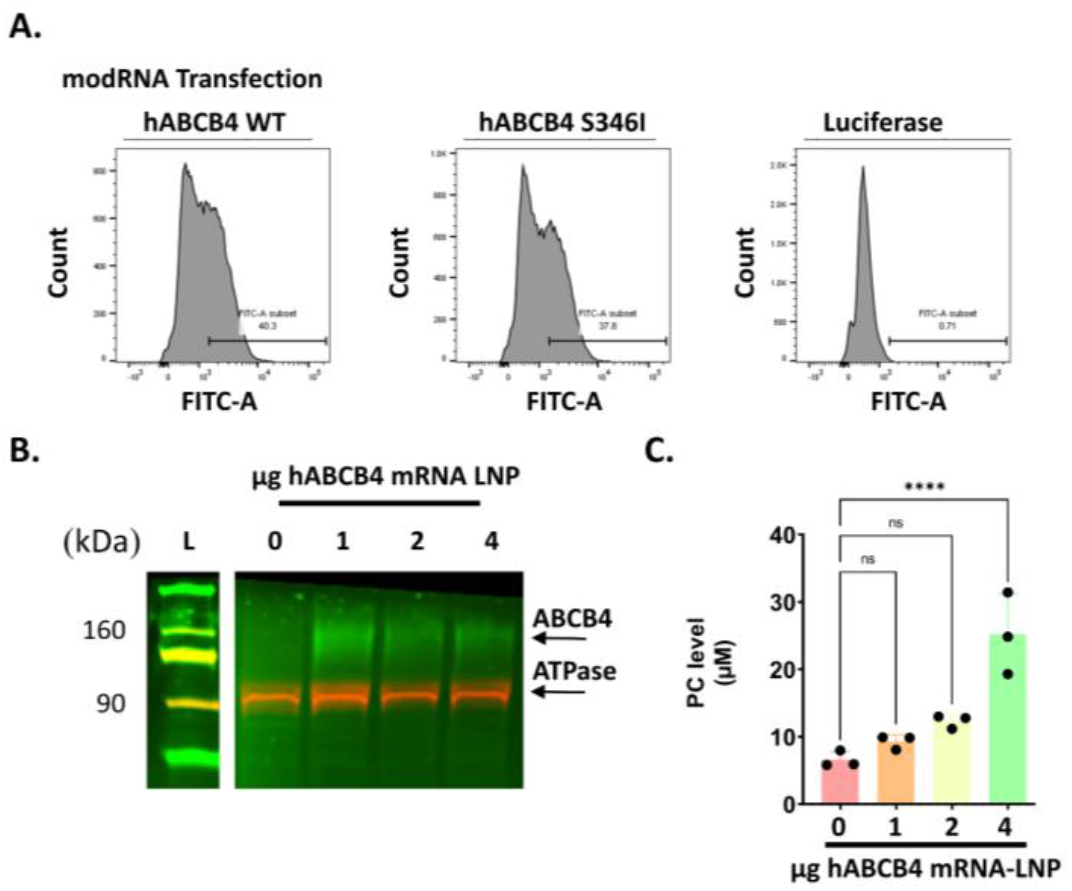
*In vitro* characterization of hABCB4 mRNA LNP. **A**. Fluorescence-activated cell sorting (FACS) analysis of Huh7 cells. hABCB4 level was significantly increased in HuH7 cells transfected with wild type and mutant hABCB4 modRNA compared to a luciferase control. **B**. Western blot analysis of Huh7 cells. hABCB4 protein level is measured in membrane fraction of HuH7 cells transfected with hABCB4 mRNA LNP, and is elevated compared to non-transfected control cells. **C**. Supernatant extracted from HuH7 cell transfected with hABCB4 mRNA LNP showed an increase in phosphatidylcholine level (PC).

### Characterization of hABCB4 mRNA LNP *in vivo*

We next examined the systemic delivery and durability of hABCB4 mRNA LNP in vivo. Wild-type FVB mice received a single injection of the hABCB4 mRNA LNP at 3 doses (0.3, 0.5, and 1 mg/kg), and livers were harvested at 6, 24, 48, and 72 hours post injections (Fig. 2A). hABCB4 mRNA expression at 1 mg/kg peaked at 6 hours post injection (1.5×10^6 copies per ug of RNA) (Fig. 2B). Next, we used an ABCB4 antibody that detects only hABCB4, and observed that hABCB4 protein expression in the liver (cell-membrane fraction) at 0.5 and 1 mg/kg was sustained up to 48 hours post injection (Fig. 2C, S1A). Furthermore, hABCB4 protein was expressed in a dose-dependent manner and expression was the highest at 1mg/kg after 48 hours compared to 0.3 and 0.5 mg/kg doses of mRNA-LNP (Fig. 2D). In addition, we used immunohistochemistry to determine the distribution of hABCB4 in the liver. hABCB4 expression was widely expressed and distributed in hepatocytes, and we observed reduced expression of hABCB4 at 72 hours post injection (Fig. 2E). Further, liver histology was examined in a blinded manner, and no hepatocellular degeneration or immune cells infiltrate were observed at any of the mRNA dose levels (data not shown). To rule out that hABCB4 mRNA LNP was delivered to tissues outside the liver, we measured the expression of hABCB4 in the spleen, kidney, and gut, and detected no expression (data not shown).

**Figure 2.**
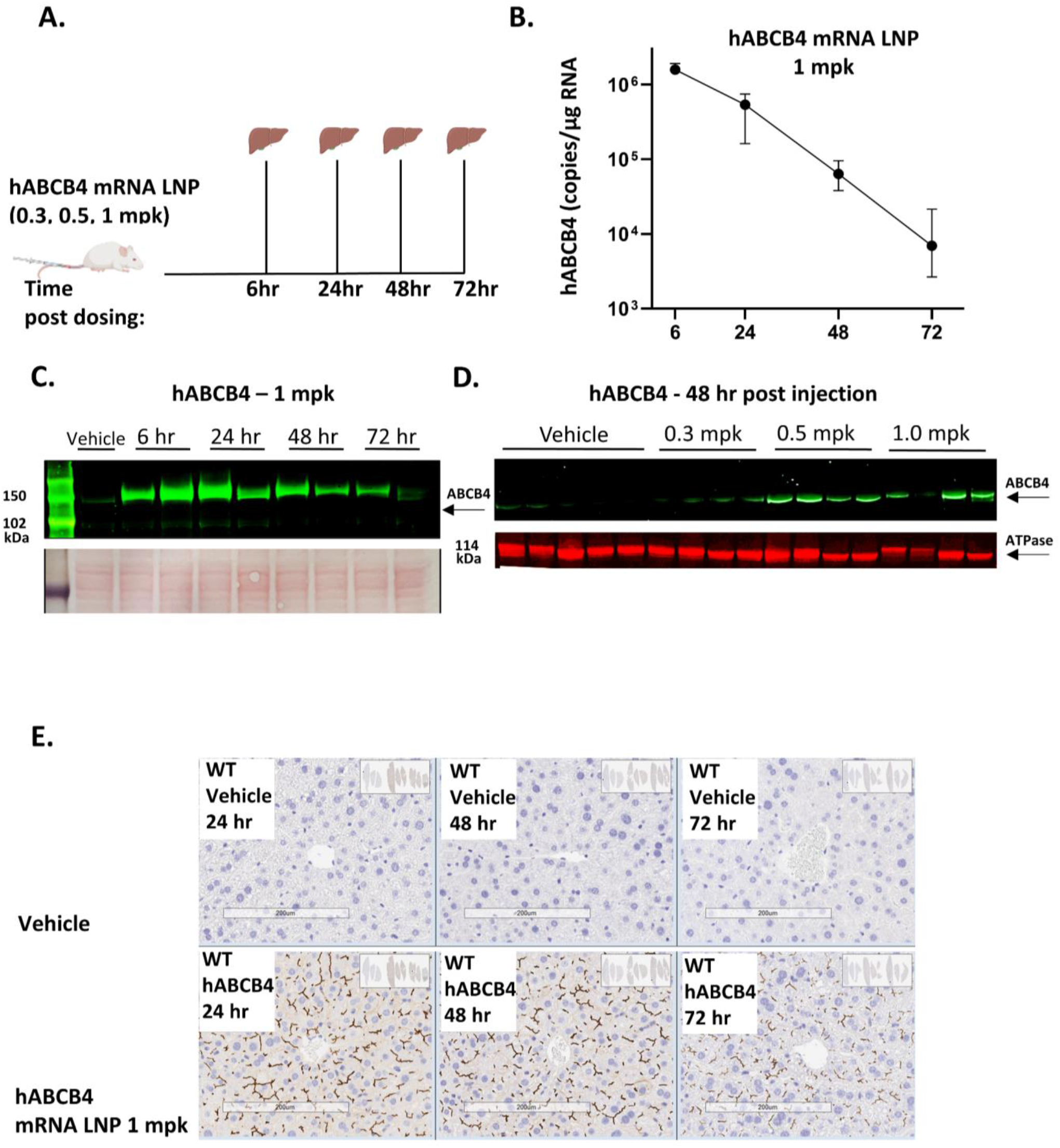
*In vivo* characterization of hABCB4 mRNA LNP in FVB wild-type mice. **A**. Schematic of the experimental design. Mice received a single injection of the hABCB4 mRNA LNP at 3 doses (0.3, 0.5, and 1 mpk), and livers were harvested at 6, 24, 48, and 72 hours post injections. **B**. Time course measurement of mRNA expression of hABCB4 in whole liver tissues was assessed using digital droplet PCR (ddPCR). **C**. hABCB4 protein level was measured in whole liver tissues (cell membrane fraction) at different harvest points using Western blotting. Ponceau whole protein stain was used as a loading control. **D**. hABCB4 and ATPase (loading control) protein levels were measured in whole liver tissues (cell membrane fraction) using different doses of hABCB4 mRNA LNP at 48 hours harvest point. **E**. Representative images of hABCB4 immunohistochemistry of vehicle treated mice (upper panel) and hABCB4 mRNA LNP treated mice (bottom panel). A significant increase in the number of hABCB4-positive cells was observed in the livers after hABCB4 mRNA LNP dosing. mpk, mg/kg.

### ABCB4 mRNA LNP restores PC level in the Abcb4^-/-^ mouse model

To measure delivery and therapeutic efficacy of hABCB4 mRNA LNP in hepatocytes of fibrotic livers *in vivo*, we generated a whole body Abcb4^-/-^ knockout mice on the FVB background. In the absence of Abcb4 protein, toxic bile without phosphatidylcholine will accumulate and lead to a cascade of events culminating with severe hepatic inflammation and fibrosis [15, 16]. 8-week-old male and female Abcb4^-/-^ mice and their counterpart wild-type controls received intravenous (i.v.) injections of 1 mg/kg hABCB4 or luciferase (control) mRNA LNP every 48 hours for two weeks - 5 doses in total - (Fig. 3A). Animals were euthanized 24 hours after the last dose, and bile, blood and tissues were collected. As expected, hABCB4 transcripts were not detected in untreated wild-type or Abcb4^-/-^ livers (Fig. 3B), but delivery of hABCB4 mRNA LNP led to a robust hABCB4 gene expression in the liver of Abcb4^-/-^ mice (Fig. 3B). Similarly, hABCB4 protein expression was not detected in untreated wild-type or Abcb4^-/-^ livers (Fig. 3C, D), and its level was significantly elevated in the livers of both female and male Abcb4^-/-^ mice after hABCB4 mRNA LNP injections (Fig. 3C, D, S1B). Furthermore, we developed a novel quantitative mass-spectrometry assay to measure both human and mouse Abcb4 protein expression in the liver. Two lead peptides were selected for quantitative assay development: (i) conserved between mouse and human ABCB4, which measures total ABCB4 when administering hABCB4 mRNA and endogenous wild-type level and (ii) human specific ABCB4 peptide that can selectively measure the human transgene protein. We observed robust hABCB4 expression (70 ng hABCB4/g liver) in the livers of Abcb4^-/-^ mice treated with hABCB4 mRNA LNP (Fig. 3E). Remarkably, human Abcb4 protein level in the Abcb4^-/-^ livers reached a similar level to the endogenous mouse Abcb4 protein level in the wild-type livers (Fig. 3F). Next, we measured phosphatidylcholine in the bile collected from the gall bladder of treated and untreated mice by mass-spectrometry. Phosphatidylcholine 16:0_18:1 and the total phosphatidylcholine pool in bile were absent in the Abcb4^-/-^ mice compared to the control group, and injections of hABCB4 mRNA LNP restored phosphatidylcholine to wild-type levels (Fig. 4A, B). These data show that LNP delivery of hABCB4 was sufficient to fully restore Abcb4 protein, and consequently, a remarkable restoration of phosphatidylcholine in bile

**Figure 3.**
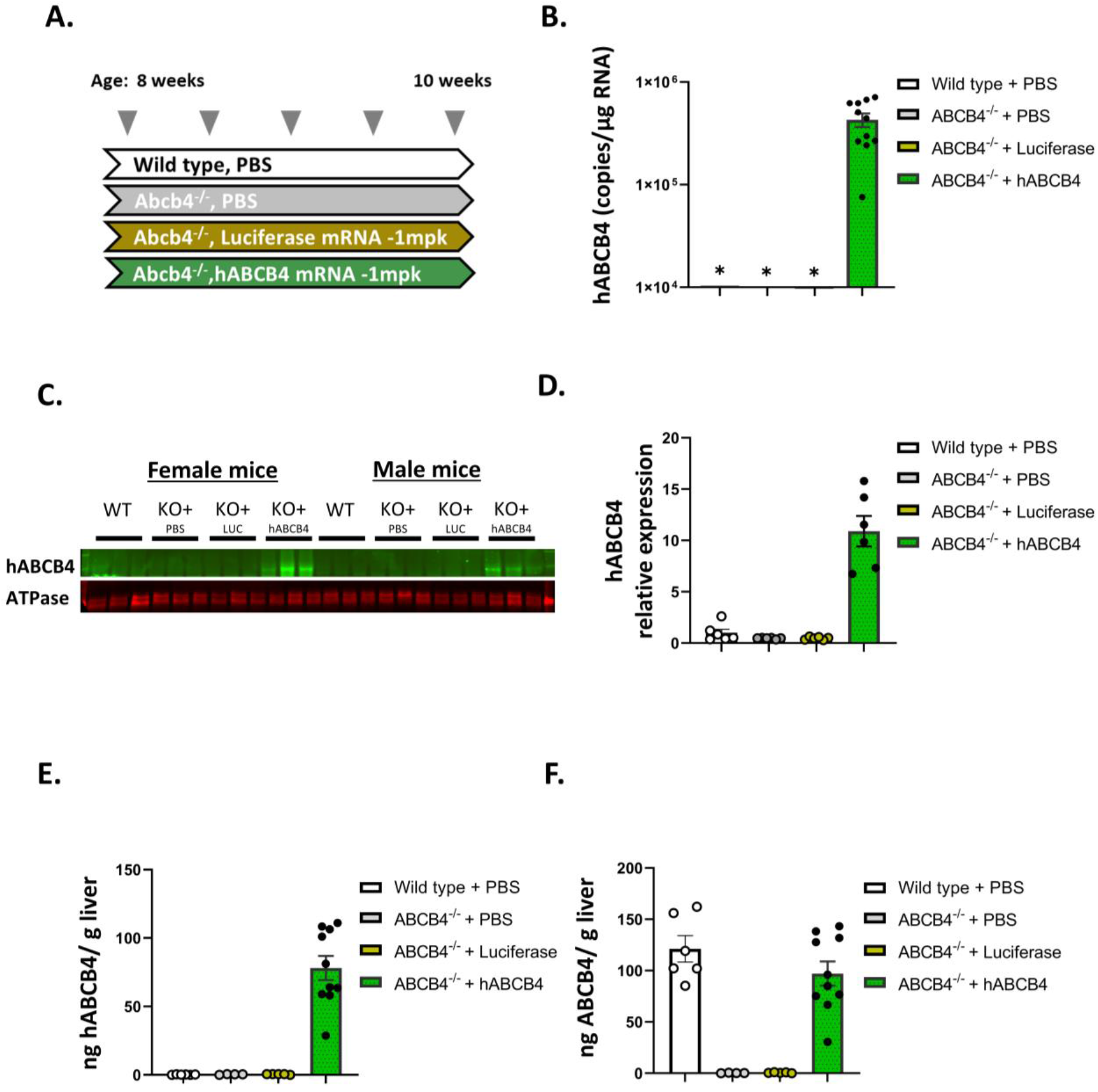
Systemic administration of hABCB4 mRNA LNP in the Abcb4^-/-^ mice. **A**. Schematic of the experimental design. Wild-type and Abcb4^-/-^ mice were dosed every 48 hours for a total of 5 doses with phosphate buffered saline (PBS), luciferase mRNA LNP, and hABCB4 mRNA LNP at 1 mpk. **B**. Gene expression analysis by ddPCR of liver homogenates. mRNA expression of hABCB4 was significantly increased in the liver of Abcb4^-/-^ mice treated with hABCB4 mRNA LNP. **C**. Western blot analysis of whole liver homogenates. hABCB4 protein level was significantly increased in whole liver tissues (cell membrane fraction) from Abcb4^-/-^ male and female mice treated with hABCB4 mRNA LNP. **D**. Quantification of hABCB4 protein level in liver lysates using Western blotting. Elevated levels of human ABCB4 protein are detected in the mouse liver following dosing with hABCB4 mRNA LNP. **E**. Novel LC:MS was developed and employed to measure hABCB4 protein level whole liver tissues (cell membrane fraction). **F**. Novel LC:MS was developed and employed to measure both hABCB4 and mouse Abcb4 protein level whole liver tissues (cell membrane fraction).

**Figure 4.**
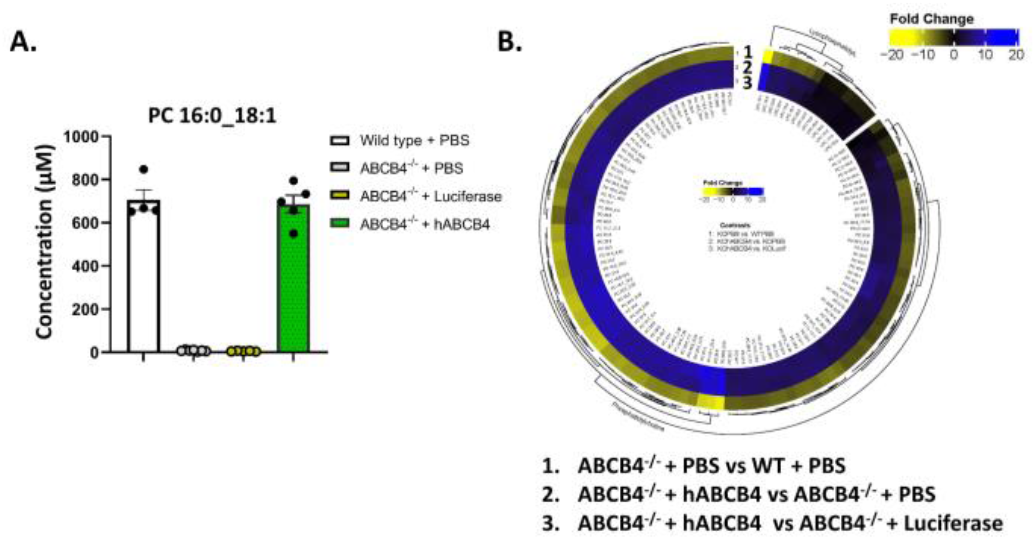
Restoration of phosphatidylcholine (PC) level in bile of Abcb4^-/-^ upon administration of hABCB4 mRNA LNP. **A**. 2-3µl of Bile was extracted from gallbladder of wildtype and Abcb4^-/-^ mice and processed for lipidomics analysis using LC:MS. PC level (16:0_18:1) was fully restored in the Abcb4^-/-^ treated with hABCB4 mRNA LNP. B. Heatmap showing absence of phosphatidylcholine profile in the bile of Abcb4^-/-^ mice, and restoration of phosphatidylcholine profile upon administration of hABCB4 mRNA LNP.

### ABCB4 mRNA LNP ameliorates liver damage and stiffness in the Abcb4^-/-^ mouse model

To address whether the restoration of Abcb4 in hepatocytes in the Abcb4^-/-^ mice suppressed the hepatocellular injury and delayed disease progression, multiple injury indices were analyzed. First, we observed that treatment with hABCB4 mRNA LNP led to a significant reduction in the mean alkaline phosphatase (ALP) level (Fig. 5A), indicating that restoration of PC level in bile reduced hepatocellular or biliary tract injury. Second, measuring liver elasticity is emerging as a non-invasive surrogate to measure liver fibrosis [17]. Thus, we employed shear wave elastography to measure liver elasticity in the Abcb4^-/-^ mice before and after hABCB4 mRNA LNP injections. As expected, the mean liver elasticity score was significantly higher in the Abcb4^-/-^ mice compared to the wild-type control (Fig. 5B). On the other hand, Abcb4^-/-^ mice treated with hABCB4 mRNA LNP had a lower and significant mean elasticity score at 10 weeks compared to baseline level at 8 weeks (Fig. 5B). Mean elasticity score was trending downward but did not reach significance between Abcb4^-/-^ mice treated with hABCB4 mRNA LNP and control groups (Abcb4^-/-^ mice treated with PBS and luciferase mRNA LNP), indicating that a longer treatment or higher concentration of hABCB4 mRNA LNP may be required to achieve a significant effect. Moreover, bulk RNA-sequencing showed that hABCB4 mRNA LNP treatment significantly decreased levels of profibrogenic and proinflammatory genes in the Abcb4^-/-^ cohort compared to the luciferase control group (Fig. 5C). Our data show for the first time that restoration of Abcb4 protein significantly decreases liver stiffness associated with the Abcb4^-/-^ mice. Furthermore, liver function enzymes and the fibrosis gene signature were down in the Abcb4^-/-^ treated mice.

**Figure 5.**
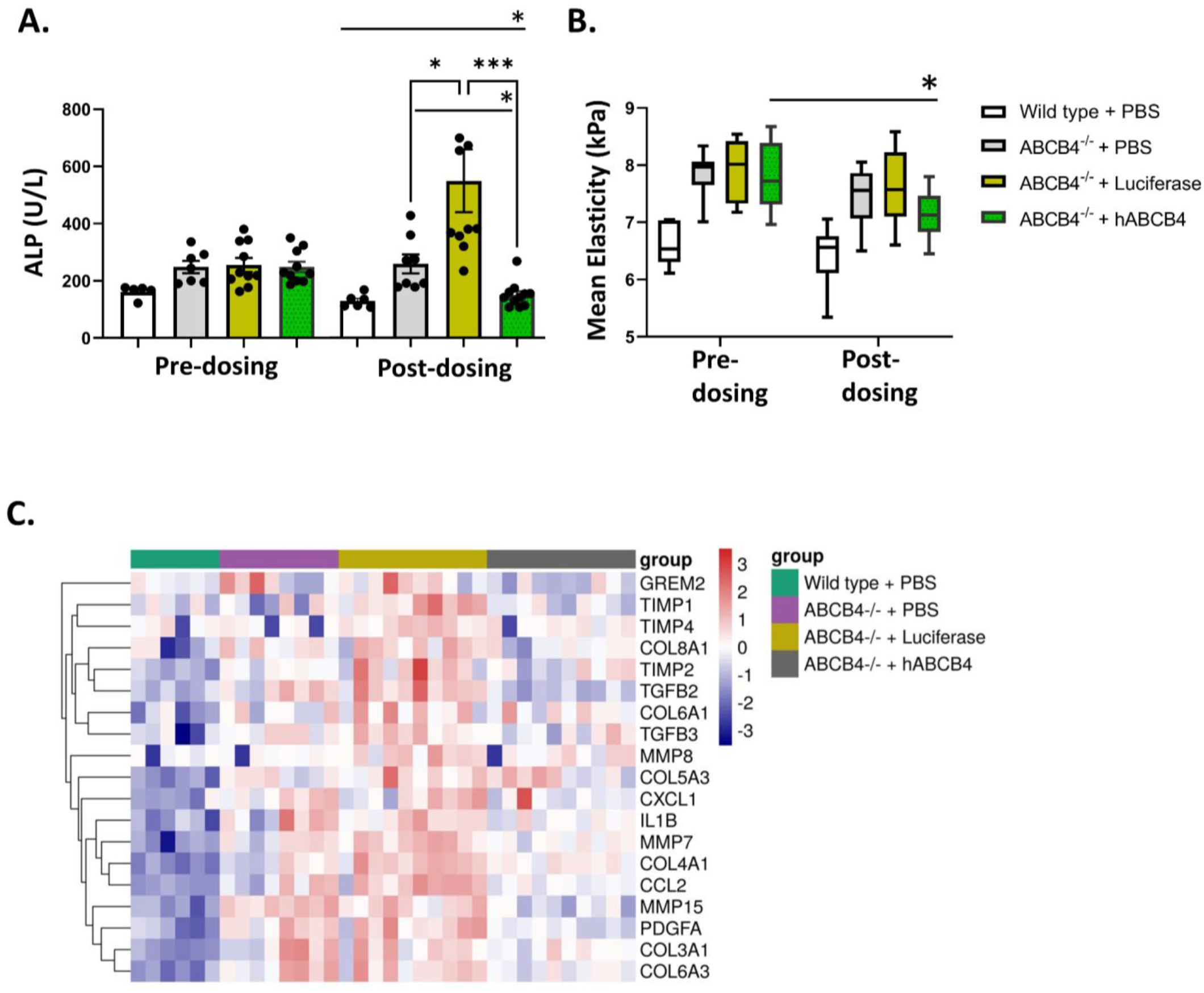
Administration of hABCB4 mRNA ameliorates liver disease phenotypes in the Abcb4^-/-^ mouse. **A**. Alkaline phosphatase (ALP) level was measured in serum of wild-type and Abcb4^-/-^ mice before and after dosing with phosphate buffered saline (PBS), luciferase, and hABCB4 mRNA LNP. Abcb4^-/-^ treated with hABCB4 mRNA exhibited a marked decrease in ALP level. **B**. Shear wave elastography was employed to measure liver stiffness in wild-type Abcb4^-/-^ mice before and after dosing with PBS, luciferase, and hABCB4 mRNA LNP. A significant decrease was observed in liver stiffness in Abcb4^-/-^ treated with hABCB4 mRNA. **C**. Heatmap showing top differentially expressed fibrotic and inflammatory genes in whole liver tissues. ALP, alkaline phosphatase.

### ABCB4 mRNA LNP ameliorates liver histopathology in the Abcb4^-/-^ mouse model

We next looked at whether short-term systemic administration of hABCB4 mRNA LNP was sufficient to impact hepatic fibrosis. First, based on histopathological qualitative evaluation of H& E stained sections, hepatic inflammation and fibrosis were scored in a blinded manner, and a significant reduction in the group treated with hABCB4 mRNA LNP was observed (Fig. 6A, B). Next, quantification of collagen level in the liver by measuring the hydroxyproline content showed a significant reduction in fibrosis in the hABCB4 mRNA LNP-treated mice compared to the Abcb4^-/-^ PBS group (Fig. 6C). Lastly, picrosirius red-stained sections were scored blindly, and a trend toward reduction in fibrosis upon hABCB4 mRNA LNP treatment was observed (Fig. 6A, D). In addition to liver fibrosis, we assessed the impact of hABCB4 expression on ductular reaction by measuring cytokeratin 19 (CK19) positive area in the liver sections. CK19 level was significantly upregulated in the Abcb4^-/-^ mice, and administration of hABCB4 mRNA led to a significant reduction compared to the control groups (Fig. 6A, E). Collectively, these findings indicate LNP delivery of hABCB4 vastly improved hepatic fibrosis in the Abcb4^-/-^ livers.

**Figure 6.**
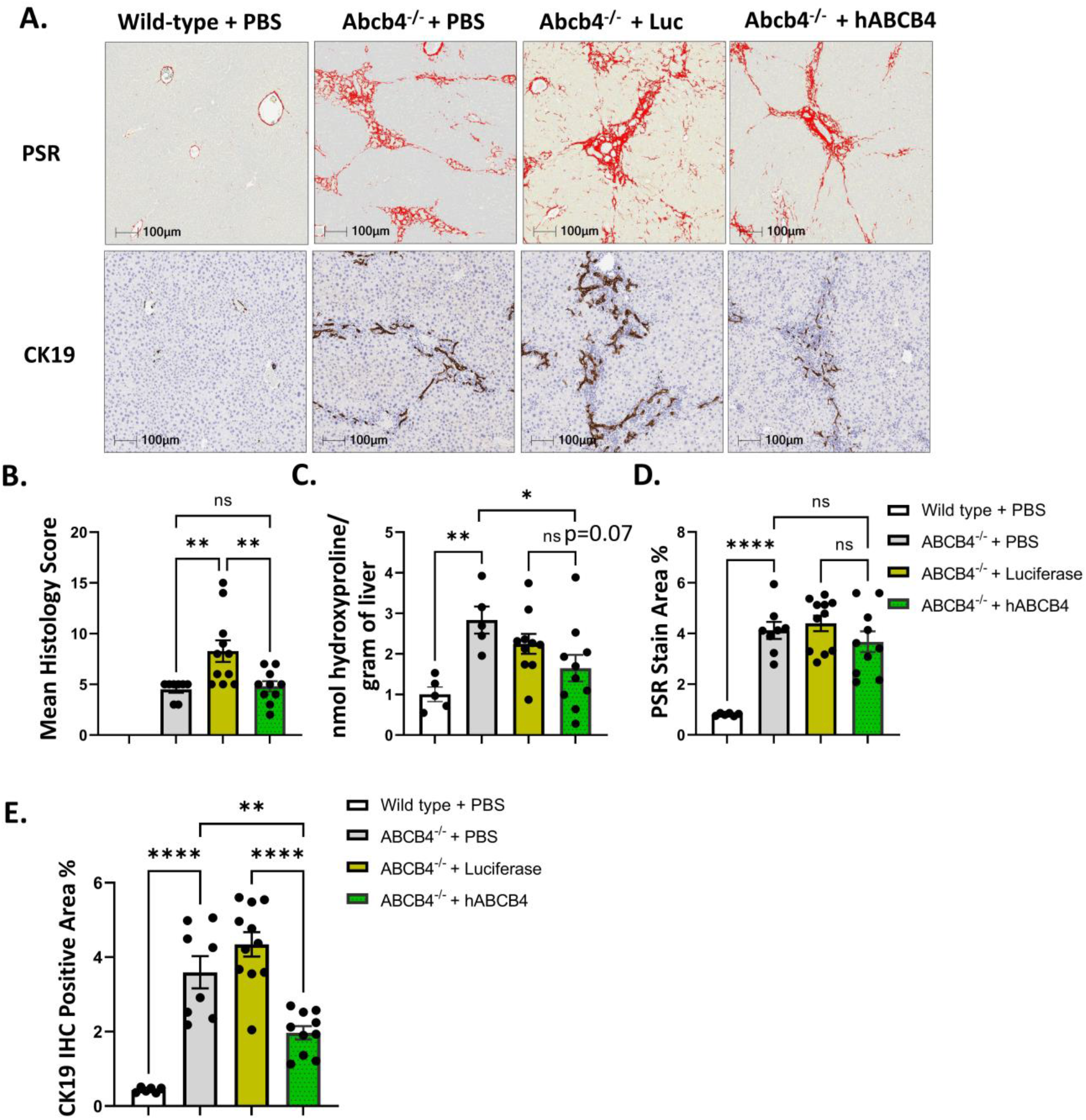
Administration of hABCB4 mRNA ameliorates fibrosis and ductular reaction in the Abcb4^-/-^ mouse. **A**. Representative images of picrosirius red (upper panel) and cytokeratin 19 (bottom panel) immunohistochemistry in wild-type and Abcb4-/mice treated with phosphate buffered saline (PBS), luciferase, and hABCB4 mRNA LNP. **B**. Qualitative quantification of mean histology score using H& E-stained liver sections. **C**. Biochemical assessment of hepatic fibrosis confirmed a markedly decreased hydroxyproline level in Abcb4^-/-^ mice treated with hABCB4 mRNA. **D**. Quantification of PSR score (shown as PSR stain area %). **E**. Quantification of CK19 positive area in liver sections. Significant reduction in CK19 observed in animals treated with hABCB5 mRNA-LNP.

In addition, these results also indicate that a longer duration of hABCB4 mRNA LNP treatment might be required to translate to significant changes across measures of fibrosis in the Abcb4^-/-^ liver.

## DISCUSSION

The major findings reported here relate to the emerging use of mRNA as a new therapeutic modality in rare liver diseases and beyond. Key findings showed that systemic delivery of hABCB4 via mRNA LNP led to: (i) a robust, durable, and evenly distributed expression of hABCB4 in the liver *in vivo* (ii) a complete restoration of PC level in bile in the Abcb4^-/-^ mice (iii) amelioration of liver stiffness and damage, and (iv) changes in liver histology including significant decreases in measures of liver fibrosis and ductular reaction.

These findings provide additional support for mRNA as a new therapeutic modality to treat rare and common diseases. Genome-wide association studies have shed light on the importance of Abcb4 in cholangiopathies including PFIC3, LPAC, and adult biliary fibrosis or cirrhosis [1, 4]. A metaanalysis of genome-wide association studies of liver enzymes from UK Biobank and Biobank Japan identified a robust genetic signal at *ABCB4* significantly associated with alanine aminotransferase (*p* = 2.0×10^−41^) and aspartate aminotransferase (*p* = 8.1×10^−11^) (Figure S2A) [18]. Abcb4 plays a key role in forming mixed lipid micelles in the bile by translocating phosphatidylcholine from the inner to the outer leaflet of the lipid bilayer of the bile canalicular membrane [2]. In patients lacking a proper functional Abcb4, toxic bile salts will accumulate leading to cholangiocytes and hepatocytes injury [1]. Over 300 disease-causing Abcb4 variants have been reported, often those with homozygous loss of the protein leading to a more devastating disease (e.g., end-stage liver disease and malignancies) and those with heterozygous loss in less severe diseases [7]. Ursodeoxycholic acid (UDCA) is currently the only drug approved by the U.S Food and Drug Administration to treat patients with cholestatic diseases (e.g., PBC, PFIC3) [19, 20]. UDCA protects liver epithelia (cholangiocytes and hepatocytes) by modifying toxic bile contents by various complex mechanisms [21]. However, there is an urgent need to find more therapies for patients with cholangiopathies that do not respond well to UDCA.

Recently, gene therapy has emerged as a revolutionary medicine for treating rare and common genetic diseases [22, 23]. Metabolic and rare liver diseases have unlimited potential for this modality since cargoes (e.g., recombinant adenoassociated virus, lipid nanoparticles) carrying genetic material to cells end up in the liver. LNPs are the most clinically advanced non-viral nucleic acid delivery system, with the successful translation of one siRNA therapeutic and two mRNA Covid-19 vaccines in the clinic [24, 25]. Systemic delivery of HNF4α, a master regulator of hepatocyte phenotype, via mRNA LNP restored gene targets and inhibited fibrosis in multiple preclinical models of liver fibrosis [14]. On the other hand, AAV-mediated gene overexpression has been widely used to treat liver fibrosis in preclinical models [26, 27]. In recent work, both AAV and mRNA LNP modalities have shown promising potential in delivering functional hABCB4 to the liver and modulating liver phenotypes in a preclinical model of PFIC3 and PSC [11, 13].

In assessing the expression and durability of hABCB4 upon systemic delivery, hABCB4 transcript peaked at 6 hours post hABCB4 mRNA administration (Fig. 2B), while the protein expression lasted for 48 hours (Fig. 2C). This short durability of mRNA and protein expression currently presents a major challenge for mRNA LNP technology compared to AAV where greater transduction efficiency and long-term efficacy in mediating gene expression have been shown in several preclinical and clinical studies [23, 28]. 1 mg/kg hABCB4 mRNA achieved a remarkable restoration of PC profile in bile to wild-type levels as measured by mass-spectrometry (Fig. 4A, B). Recent publication by Wei et al also demonstrated that repeated injections of hABCB4 led to 42% restoration of PC levels in bile [11]. As expected, PC restoration led to amelioration of various liver phenotypes including liver stiffness as measured by shear wave elastography (Fig. 5B). To our knowledge, this has been the first time shear wave elastography technology is used in the Abcb4^-/-^ pre-clinical model of PFIC3 and PSC to predict resolution of fibrosis. Our data further showed that liver fibrosis, as scored by histology evaluation or hydroxyproline, was significantly ameliorated in the Abcb4^-/-^ hABCB4 group (Fig. 6A, D). Similarly, Wei et al showed that two-weeks administration of hABCB4 halted

## ACKNOWLEDGMENT

The authors would like to thank the BioMedicine Design (BMD) team (Jin Li, Ruiting Lin, Linette Rodriguez, Chong Wang, Divya Patel, Luke Luo, Martina Zafferani, Aaron D’Antona, Justin Cohen, Amy Tam, and Laura Lin) for the preparation, generation, and QC of the hABCB4 mRNA-LNP; James Stejskal and Deborah Burt for their help running liver biomarker analysis; Dominica Lopes and the Comparative Medicine team for their efforts with the *in vivo* studies; David Morrisey, Uwe Schoenbeck, Morten Sogaard, and Pfizer colleagues in Emerging Science and Innovation for their support and stimulating discussion. We are grateful to Greg LaRosa and Robert Moccia for discussion and insights and on genetics and disease indication.

## AUTHOR CONTRIBUTION

Experiments were performed by the following authors: YW, YC, and MH (vectors design and generation), YA (mice generation), GYL, CC and MA (in vitro PC assay & WB), RK and XS (histopathology analyses), ZJ and JH (ddPCR), MA, BL and ND (bile extraction and LC-MS), AH and DH (SWE), RO (LCprogression of liver fibrosis as measured by PSR and hydroxyproline content. Future studies are warranted to determine what level of PC restoration is required to achieve a therapeutic index to ameliorate liver phenotypes associated with loss of Abcb4. It is also worth noting that a significant increase in ALP and AST levels were observed in the luciferase mRNA LNP group compared to the PBS control group (a similar observation in few fibrotic and inflammatory genes). Future studies in preclinical models need to evaluate proper controls for mRNA studies (e.g., different reporter genes, mRNAs of different nucleotide compositions and/or UTRs).

In conclusion, our data continue to support an important pathophysiological role of Abcb4 in rare biliary disease, and suggest the mRNA-LNP modality as an attractive therapeutic approach for rare and common liver diseases. The current work suggests that administrating hABCB4 mRNA is safe in wildtype and genetically modified mice and has the potential to be translated to the clinic to treat patients with Abcb4 mutations. In the future, investigations should determine the therapeutic benefit of overexpressing hABCB4 beyond cholestatic diseases. One could investigate whether administration of hABCB4 mRNA could ameliorate liver fibrosis in pre-clinical model of alcoholic and non-alcoholic liver diseases.

MS), SL (bulk RNA-seq analysis). BL and MA led and executed all *in vivo* studies. SHC, MA, MH, and MS contributed to conception, design, and/or interpretation of the work. MA wrote the manuscript with input from all authors. MTS; MS; MH helped with revision and drafting of manuscript.

This work was supported by Pfizer.

