## Supplemental Figures 1 and 2 for "Targeting ABCB4 using mRNA-LNP for the treatment of rare liver diseases"

#### Supplementary Figure 1:

**A.**

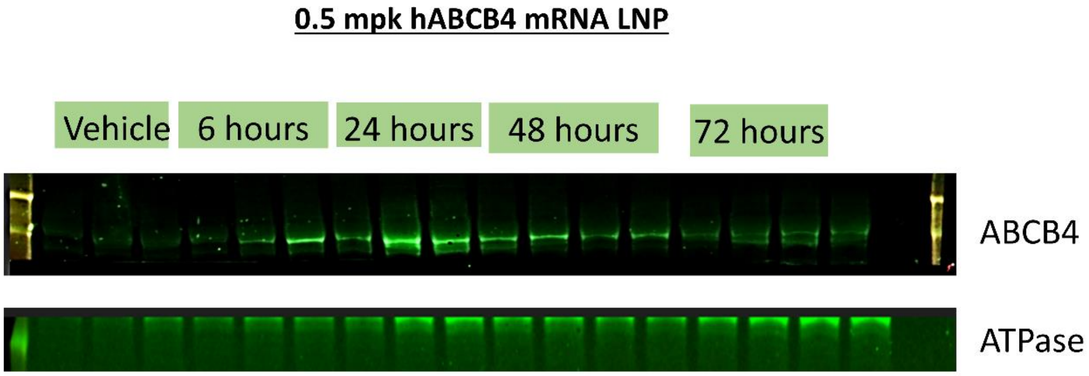


**B.**

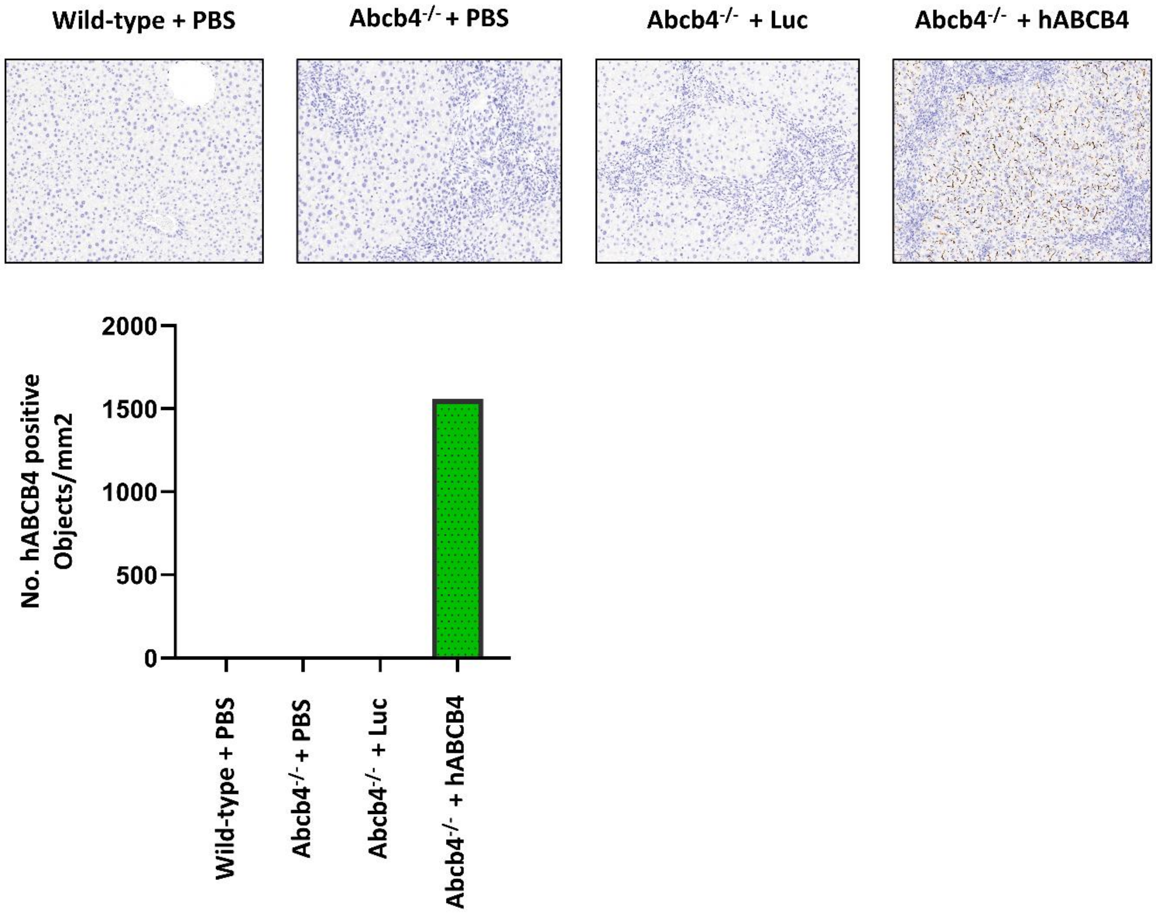

**Supplementary Figure 1. Administration of hABCB4 mRNA leads to a widely distributed and durable expression of hABCB4 protein. A.** hABCB4 protein level was measured in whole liver tissues (cell membrane fraction) from wild-type mice injected with 0.5 mpk hABCB4 mRNA LNP at different harvest points. **B**. Representative images of hABCB4 immunohistochemistry of PBS treated wild-type mice, PBS treated Abcb4$^{-/-}$ mice, luciferase mRNA LNP treated Abcb4$^{-/-}$ mice, and hABCB4 mRNA LNP treated Abcb4$^{-/-}$ mice. A significant increase in the number of hABCB4-positive cells was observed in the livers after hABCB4 mRNA LNP dosing. Mpk, mg/kg; PBS, phosphate buffered saline.

### Supplementary Figure 2:

## A.

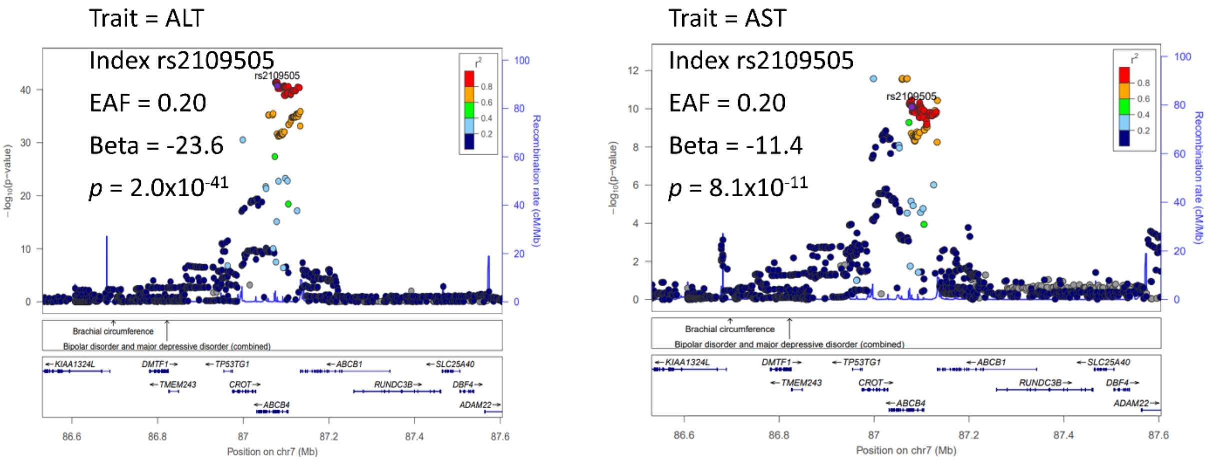

**Supplementary Figure 2. Robust association of ABCB4 locus with liver enzyme levels. A.** Meta-analysis of genome-wide association studies of liver enzymes from UK Biobank (UKBB) and BioBank Japan (BBJ) identified a robust genetic signal at *ABCB4* significantly associated with alanine aminotransferase (ALT, $p = 2.0\text{x}10^{-41}$) and aspartate aminotransferase (AST, $p = 8.1\text{x}10^{-11}$).
